# Developmental Toxicity and Lethality of Structurally Diverse PFAS in Zebrafish

**DOI:** 10.1101/2025.08.07.669106

**Authors:** Matthew Farrell, Ria Bakshi, Emily Griffith, Antonio Planchart

## Abstract

Per-and polyfluoroalkyl substances (PFAS) are ubiquitous environmental contaminants that have been associated with adverse health effects in highly exposed populations. Manufacturers have taken steps to replace toxic long-chain perfluoroalkyl acids (PFAAs) with short-chain PFAAs and perfluoroether acids (PFEAs). There is little to no toxicity data for many of these chemicals. Most of the data that are available are taken from studies that do not account for the pH of highly concentrated PFAS solutions, resulting in highly acidic conditions that do not accurately reflect real-world exposures. The goal of this study was to evaluate the lethality and developmental toxicity of 17 structurally diverse PFAS in zebrafish in a pH-neutral environment. We then compared results to determine the impacts of chain length, head group, and ether linkages on toxicity. The potency of PFAS to induce mortality and developmental toxicity endpoints increased with chain length, and sulfonic acids were more potent than carboxylic acids. The inclusion of ether oxygens was associated with reduced potency relative to PFAAs with equal chain length and head group. Perfluorooctane sulfonic acid (PFOS) was the most potent compound, followed by perfluoroundecanoic acid (PFUnDA) and perfluorododecanoic acid (PFDoDA). Short-chain compounds perfluoro-2-methoxyacetic acid (PFMOAA) and perfluorobutanoic acid (PFBA) were the least potent. These data were used to construct a multiple linear regression model for PFAS potency. Failed swim bladder inflation was the most sensitive developmental toxicity endpoint for all assessed chemicals. Other common phenotypes included spinal curvature, edema, craniofacial malformations, and ocular malformations. Ocular malformations were more common in response to sulfonic acids. No other phenotypes exhibited significant structural specificity. Our study provides new toxicity data for a diverse set of PFAS under environmentally relevant conditions. Future studies should be expanded to include more branched structures and head groups not present in our testing set to allow for improved understanding of how other structural features impact toxicity.

## INTRODUCTION

Per- and polyfluoroalkyl substances (PFAS) are synthetic chemicals first produced in the late 1930s (Gaines, 2023). These fluorinated compounds are excellent surfactants and are extremely stable, making them useful in chemical manufacturing. PFAS are both hydrophobic and lipophobic, which has led to their use in many consumer products. PFAS can be found in textiles, electronics, nonstick cookware, firefighting foams, and many other products (Glüge et al., 2020). In these cases, PFAS frequently serve as water- or stain-resistant coatings. PFAS are a constantly growing chemical class, with more than 14,000 structures currently listed in the US Environmental Protection Agency’s CompTox Chemicals Dashboard (Williams et al., 2017). New PFAS are produced both intentionally and as incidental byproducts in manufacturing processes.

The highly stable carbon-fluorine bond is key to the performance of PFAS in their many roles. However, this stability is also a major factor in the growing concern over PFAS as environmental contaminants. PFAS are highly resistant to environmental degradation and metabolic transformation (Cousins et al., 2020). PFAS can be released into the environment either directly, from point sources like fluorochemical plants and aqueous film-forming foam (AFFF) usage, or indirectly, via landfill leachate and wastewater treatment plants (Dasu et al., 2022). Once in the environment, many PFAS are mobile and can be carried great distances through surface and groundwater, marine currents, and the atmosphere (Ahrens et al., 2023). As a result of this mobility and persistence, PFAS are globally ubiquitous contaminants. They are present at measurable levels in soil, water, and the atmosphere around the globe (Panieri et al., 2022). Multiple large-scale cohort studies have detected PFAS in the serum of nearly every person tested (Calafat et al., 2007; Forns et al., 2020; Tian et al., 2018). These contaminants have been found at high levels even in arctic wildlife, far from known sources (Giesy and Kannan, 2001).

PFAS have been connected to numerous adverse health outcomes such as dyslipidemia, immunosuppression, thyroid disease, and cancer (NASEM et al., 2022). PFAS are also developmental toxicants. Maternal PFAS are readily transferred via the placenta and breast milk, resulting in reduced birth weights and weaker vaccine responses in human infants (Fenton et al., 2021). These and other developmental toxicity outcomes have been consistently replicated in animal models (Truong et al., 2022; Conley et al., 2021). However, the vast majority of health studies involving PFAS have focused on just a handful of legacy compounds, most notably perfluorooctanoic acid (PFOA) and perfluorooctane sulfonic acid (PFOS). In response to the growing concern over these chemicals, fluorochemical manufacturers have largely replaced production of these older compounds with PFEAs and short-chain alternatives. These PFAS now constitute a major part of exposure for most individuals, and many of these chemicals have little or no toxicity data (Sun et al., 2016). This gap in knowledge represents an urgent priority, as remediation of these persistent chemicals is very difficult.

The zebrafish is a well-established toxicological model that presents a number of advantages for studying PFAS. Zebrafish produce hundreds of embryos from single spawning events, and a short life cycle means that they can reach reproductive maturity as early as three months (Parichy et al., 2009). These factors combined with their low space requirements allow zebrafish to be used for high-throughput assays that are not possible with traditional mammalian models. While the environment of the adult zebrafish differs significantly from that of humans, aqueous exposures in zebrafish embryos mimic *in utero* exposures during early human development (Bugel et al., 2014). Additionally, zebrafish are transparent during early stages, facilitating observation of their internal development.

We leveraged the zebrafish model to address the paucity of PFAS toxicity data by examining the lethality and developmental toxicity of 17 different PFAS. We included a range of structures to allow identification of factors that determine potency, such as chain length, head group, and the inclusion of ether oxygens.

## METHODS

### Chemical list and sources

Perfluorododecanoic acid (PFDoDA; CAS 307-55-1), perfluoroundecanoic acid (PFUnDA; CAS 2058-94-8), perfluorodecanoic acid (PFDA; CAS 335-76-2), perfluoro-n-octanoic acid (PFOA; CAS 335-67-1), perfluoroheptanoic acid (PFHpA; CAS 375-85-9), perfluoropentanoic acid (PFPeA; CAS 2706-90-3), heptafluorobutyric acid (PFBA; CAS 375-22-4), perfluorooctane sulfonic acid (PFOS; CAS 1763-23-1), perfluoropentane sulfonic acid (PFPeS; CAS 2706-91-4), nonafluorobutane sulfonic acid (PFBS; CAS 375-73-5), undecafluoro-2-methyl-3-oxahexanoic acid (HFPO-DA (GenX); CAS 13252-13-6), and 7H-perfluoro-4-methyl-3,6-dioxaoctanesulfonic acid (Nafion byproduct 2 (NBP2); CAS 749836-20-2) were purchased from Synquest Laboratories (Alachua, FL, USA). Perfluorononanoic acid (PFNA; CAS 375-95-1), undecafluorohexanoic acid (PFHxA; CAS 307-24-4), and tridecafluorohexane-1-sulfonic acid potassium salt (PFHxS; CAS 3871-99-6) were purchased from Sigma-Aldrich (St. Louis, MO, USA). Sodium 2,2-difluoro-2-(trifluoromethoxy)acetate (PFMOAA; CAS 21837-98-9) and sodium undecafluoro-2, 4, 6, 8-tetraoxadecan-10-oate (PFO4DA; CAS 1035377-21-9) were purchased from Fluoryx Labs (Carson City, NV, USA).

### Zebrafish husbandry

A wild type zebrafish line was derived from AB wild type zebrafish obtained from the Zebrafish International Resource Center (ZIRC). Zebrafish were maintained in a carbon-filtered aquarium system at 28±1°C on a 14:10 hour day/night cycle. Tanks were kept at a density of 10 fish per liter or less. Salinity and pH were maintained at 800±50 µS and 7.5±0.5, respectively, via the addition of Instant Ocean salts and sodium bicarbonate. Embryos were stored in an incubator at 28.5°C with a 14:10 hour day/night cycle for the duration of the experiments. All embryos were placed in 0.5X E2 embryo media, consisting of 15mM NaCl, 0.5 mM KCl, 1 mM MgSO_4_, 0.15 mM KH_2_PO_4_, 0.05 mM Na_2_HPO_4_, and 0.7 mM NaHCO_3_. The NC State University Institutional Animal Care and Use Committee (IACUC, #24-231) reviewed and approved all procedures and uses of zebrafish.

### PFAS exposures

Five-day exposures of zebrafish embryos to PFAS were performed to evaluate developmental toxicity and mortality. Exposures were carried out in polystyrene six-well plates (Genessee Scientific, cat #:25-105) (Figure 1). Each well was filled with 5mL of 0.5X E2 growth media containing the target concentration of an individual PFAS. For every PFAS, 11 different concentrations and a negative control were tested across 2 well plates. Exposure ranges were individually selected for each chemical (Table 1) based on pilot studies and existing literature with the intent of capturing the full range from no observed adverse effects to 100% mortality. Exposure concentrations increased by quarter-log increments within each replicate. In the case of PFDA, PFNA, and PFHxS, the mortality increase between quarter-log concentrations was too large to accurately estimate LC_50_ values. For those PFAS, additional trials with eighth-log increments in concentration were performed to improve resolution. Most exposure solutions were prepared from PFAS stock solutions in DMSO. Perfluoroether acids (PFEAs) that are not stable in DMSO were made as aqueous stock solutions (Zhang et al., 2022). For non-ether PFAS, the final DMSO concentration in all wells was normalized to 0.8% v/v. DMSO was omitted from all PFEA exposures to ensure consistency between them. Any solutions with pH lower than the 0.5X E2 embryo media (∼7.3) were titrated with 1M NaHCO_3_ until they reached that point.

**Figure 1:**
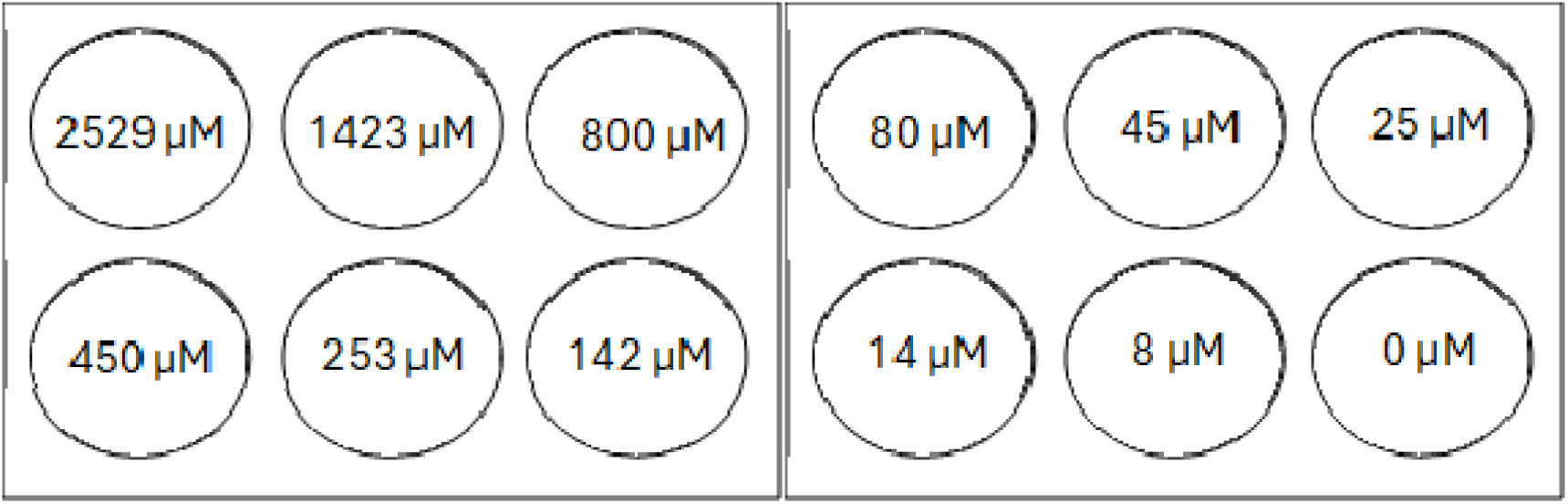
PFAS exposure setup, using PFOA concentrations as an example. Each replicate consisted of two six-well plates with exposure concentrations increasing in quarter-log increments. Each well contained 5 mL of exposure solution and 20 zebrafish embryos.

**Table 1.**
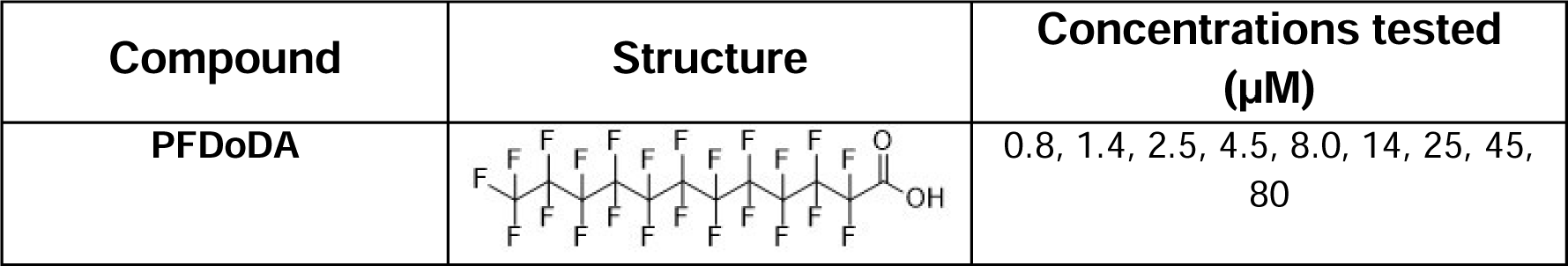
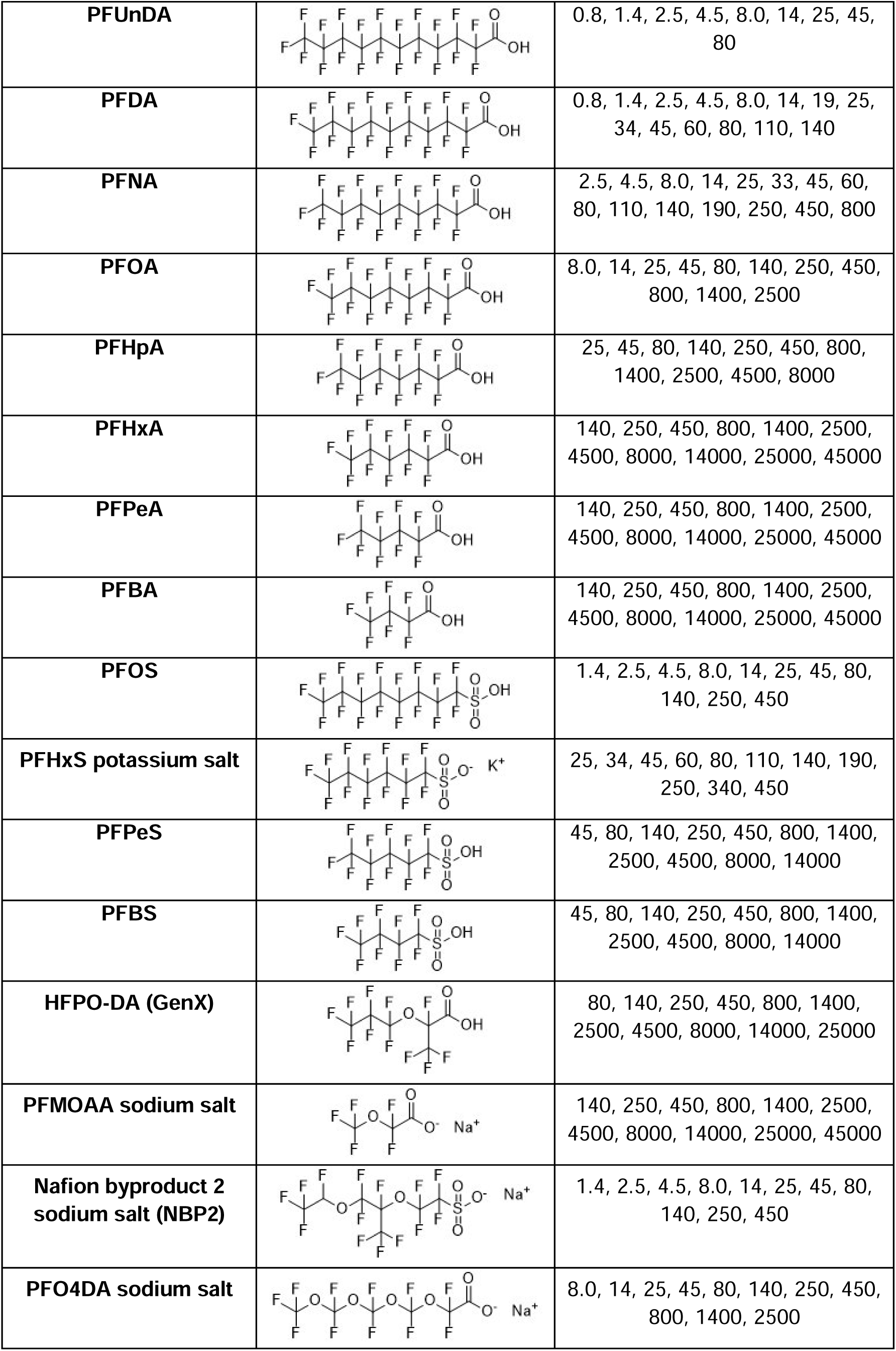
Exposure ranges and structures for all PFAS. 45,000 µM was set as an absolute maximum to prevent excess waste of PFAS with low potency. PFDoDA and PFUnDA were not fully soluble above 80 µM, and so only nine total concentrations were tested for these compounds.

| Compound | Structure | Concentrations tested (µM) |
| --- | --- | --- |
| PFDODA |  | 0.8, 1.4, 2.5, 4.5, 8.0, 14, 25, 45, 80 |
| **PFUnDA** |  | 0.8, 1.4, 2.5, 4.5, 8.0, 14, 25, 45, 80 |
| **PFDA** |  | 0.8, 1.4, 2.5, 4.5, 8.0, 14, 19, 25, 34, 45, 60, 80, 110, 140 |
| **PFNA** |  | 2.5, 4.5, 8.0, 14, 25, 33, 45, 60, 80, 110, 140, 190, 250, 450, 800 |
| **PFOA** |  | 8.0, 14, 25, 45, 80, 140, 250, 450, 800, 1400, 2500 |
| **PFHpA** |  | 25, 45, 80, 140, 250, 450, 800, 1400, 2500, 4500, 8000 |
| **PFHxA** |  | 140, 250, 450, 800, 1400, 2500, 4500, 8000, 14000, 25000, 45000 |
| **PFPeA** |  | 140, 250, 450, 800, 1400, 2500, 4500, 8000, 14000, 25000, 45000 |
| **PFBA** |  | 140, 250, 450, 800, 1400, 2500, 4500, 8000, 14000, 25000, 45000 |
| **PFOS** |  | 1.4, 2.5, 4.5, 8.0, 14, 25, 45, 80, 140, 250, 450 |
| **PFHxS potassium salt** |  | 25, 34, 45, 60, 80, 110, 140, 190, 250, 340, 450 |
| **PFPeS** |  | 45, 80, 140, 250, 450, 800, 1400, 2500, 4500, 8000, 14000 |
| **PFBS** |  | 45, 80, 140, 250, 450, 800, 1400, 2500, 4500, 8000, 14000 |
| **HFPO-DA (GenX)** |  | 80, 140, 250, 450, 800, 1400, 2500, 4500, 8000, 14000, 25000 |
| **PFMOAA sodium salt** |  | 140, 250, 450, 800, 1400, 2500, 4500, 8000, 14000, 25000, 45000 |
| **Nafion byproduct 2 sodium salt (NBP2)** |  | 1.4, 2.5, 4.5, 8.0, 14, 25, 45, 80, 140, 250, 450 |
| **PFO4DA sodium salt** |  | 8.0, 14, 25, 45, 80, 140, 250, 450, 800, 1400, 2500 |

20 wild type zebrafish embryos were placed into each well at three hours post-fertilization (hpf), corresponding to the onset of gastrulation, to minimize non-specific developmental abnormalities due to poor egg quality. Well plates were covered with parafilm to minimize evaporation and moved into an incubator. Embryos were checked daily, and any dead embryos were removed to preserve water quality. A minimum of three replicate trials were performed for every tested chemical.

### Toxicity assessments

Mortality in all wells was recorded at 24 hpf and every 24 hours thereafter. At 120 hpf, developmental toxicity phenotypes were assessed in all surviving larvae (the larval stage begins around 2-3 days). Larvae were examined for failed swim bladder inflation, spinal curvature, pericardial and yolk sac edema, craniofacial malformations, ocular malformations, and delayed hatching. These endpoints were selected based on observations in pilot studies and commonly reported phenotypes in the literature. All reported developmental toxicity phenotypes were visually identified using a dissecting microscope. Replicates with 50% or higher mortality in negative control wells were excluded from analysis.

LC_50_ and EC_50_ of failed swim bladder inflation were used as measurements of lethality and developmental toxicity for the creation of a multiple linear regression (MLR) model that predicted PFAS potency based on structural characteristics. Chain length, head group, and the number of ether oxygens were assessed for their impact on toxicity. Following literature precedent, chain length was defined as the sum of carbon (including branches), sulfur, and ether oxygen atoms (Cheng et al., 2025).

### Statistical analysis

Proportional mortality of the larvae in each experimental well was calculated after subtracting out background mortality from 0.8% DMSO wells. Any negative values generated by this adjustment were set to 0. These proportions were used to determine LC_50_, LC_20_, and LC_10_ values through nonlinear regression with a Hill slope curve fitting. The reported 95% confidence intervals (CIs) for these values are profile-likelihood CIs. Curve fitting and CI calculations were performed using GraphPad Prism 10.5.0.

For the developmental toxicity assays, each endpoint was assessed as the proportion of surviving larvae presenting each phenotype. Wells with fewer than five surviving larvae were excluded from further analysis. The incidence of each developmental toxicity phenotype in negative control wells was used as a baseline and subtracted out from experimental wells. The control-adjusted proportions were used to estimate EC_50,_ EC_20_, and EC_10_ values for each endpoint as described for the mortality assessment.

Overall frequency of each phenotype was determined by summing the affected larvae across all experimental wells and subtracting out the expected count based on extrapolation from the proportions in 0.8% DMSO negative control wells.

A multiple linear regression (MLR) model was used to correlate LC_50_ and EC_50_ values with PFAS structural properties. Chain length, head group, and number of oxygens were used as variables for our model due to their previously established importance to PFAS toxicity and binding (Cheng et al., 2025; Patlewicz et al., 2022). This MLR model was created using GraphPad Prism 10.5.0.

## RESULTS

### Lethality and potency of PFAS in developing zebrafish

15 of the 17 tested PFAS caused elevated mortality in a dose-dependent manner (Table 2). PFOS was the most potent compound, with an LC_50_ [95%CI] of 3.85 [3.31-4.46] µM. The LC_50_ of PFOS was the lowest by an order of magnitude, followed by PFUnDA and PFDoDA at 15.88 [14.0-18.1] µM and 26.32 [24.4-28.3] µM, respectively. PFBA and PFPeA had the lowest potencies of all tested compounds. For most compounds, the order of potency was unchanged whether evaluated using LC_50_, LC_20_, or LC_10_. PFPeA and GenX are notable exceptions, as both were relatively more potent when ranked using LC_10_. Both compounds, particularly PFPeA, had broad dose-response curves compared to other tested PFAS. PFMOAA and PFO4DA did not cause elevated mortality even at the highest tested concentrations (45 mM PFMOAA, 2500 µM PFO4DA).

**Table 2.** Mortality of zebrafish larvae 120 hours post-fertilization following exposure to the specified PFAS. Mortality is represented through median lethal concentration (LC_50_), 20% mortality (LC_20_), and 10% mortality (LC_10_). Some compounds did not have sufficient mid-curve experimental data to accurately calculate CIs, denoted with a ‘?’. Compounds which did not exhibit dose-dependent mortality at any tested concentration are labeled N/A. PFCA: perfluorocarboxylic acid, PFSA: perfluorosulfonic acid, PFECA: perfluoroethercarboxylic acid, PFESA: perfluoroethersulfonic acid.

|  | Compound | LC<sub>50</sub> [95% CI]<br>(μM) | LC<sub>20</sub> [95% CI]<br>(μM) | LC<sub>10</sub> [95% CI]<br>(μM) |
| --- | --- | --- | --- | --- |
| **PFCAs** | **PFDoDA** | 26.32 [24.36-28.27] | 19.84 [16.89-?] | 16.82 [13.43-?] |
|  | **PFUnDA** | 15.88 [14.02-18.09] | 11.8 [8.95-13.88] | 9.92 [6.68-?] |
|  | **PFDA** | 48.86 [45.49-51.85] | 41.17 [33.44-45] | 37.25 [27.62-?] |
|  | **PFNA** | 145.9 [126.6-167.9] | 95.42 [72.3-117.5] | 74.44 [50.01-98.89] |
|  | **PFOA** | 1045 [902.5-1204] | 655.3 [494.3-817] | 498.7 [335.7-668.8] |
|  | **PFHpA** | 6075 [5637-6576] | 5023 [4620-5662] | 4495 [4042-5203] |
|  | **PFHxA** | 12374 [11392-?] | 9860 [8528-?] | 8634 [7095-?] |
|  | **PFPeA** | 31955 [20771-79948] | 8139 [2746-15935] | 3657 [540.3-10273] |
|  | **PFBA** | 38130 [28070-75738] | 17402 [9068-26791] | 10999 [3501-20765] |
| **PFSAs** | **PFOS** | 3.85 [3.31-4.46] | 2.39 [1.82-3.02] | 1.81 [1.24-2.5] |
|  | **PFHxS** | 128.5 [121.3-?] | 120.4 [111.4-?] | 115.8 [105.4-?] |
|  | **PFPeS** | 764.3 [577.2-997.5] | 451.0 [281.5-687.7] | 331.2 [173.7-588.3] |
|  | **PFBS** | 3842 [2733-5452] | 1572 [898.8-2508] | 932.2 [421-1723] |
| **PFECA**<br>**s** | **HFPO-DA**<br>**(GenX)** | 7437 [4944-10568] | 4214 [1309-?] | 3023 [541.6-?] |
|  | **PFMOAA** | N/A | N/A | N/A |
|  | **PFO4DA** | N/A | N/A | N/A |
| **PFESA**<br>**s** | **Nafion**<br>**Byproduct 2** | 54.24 [47.03-62.76] | 35.63 [27.9-43.33] | 27.87 [19.71-36.06] |

### Developmental toxicity and potency of PFAS in zebrafish larvae

Developmental toxicity was observed for all 17 PFAS within the tested concentration ranges. Failed swim bladder inflation was the most sensitive and most common phenotype across all compounds (Table 3). The ordering of potency as a developmental toxicant was nearly identical when compared with mortality. 14 PFAS showed a sufficiently strong dose-response relationship to derive EC_50_ values for failed swim bladder inflation. In all cases, the EC_50_ was within one order of magnitude of the LC_50_ for the same compound. PFPeA, PFBA, and PFMOAA did not exhibit a dose-dependent increase in failed swim bladder inflation within the tested exposure range. However, total incidence of uninflated swim bladders was significantly increased for all three PFAS (Figure 3). No significant difference in hatching time was observed between exposed and control fish for any compound.

**Figure 2.**
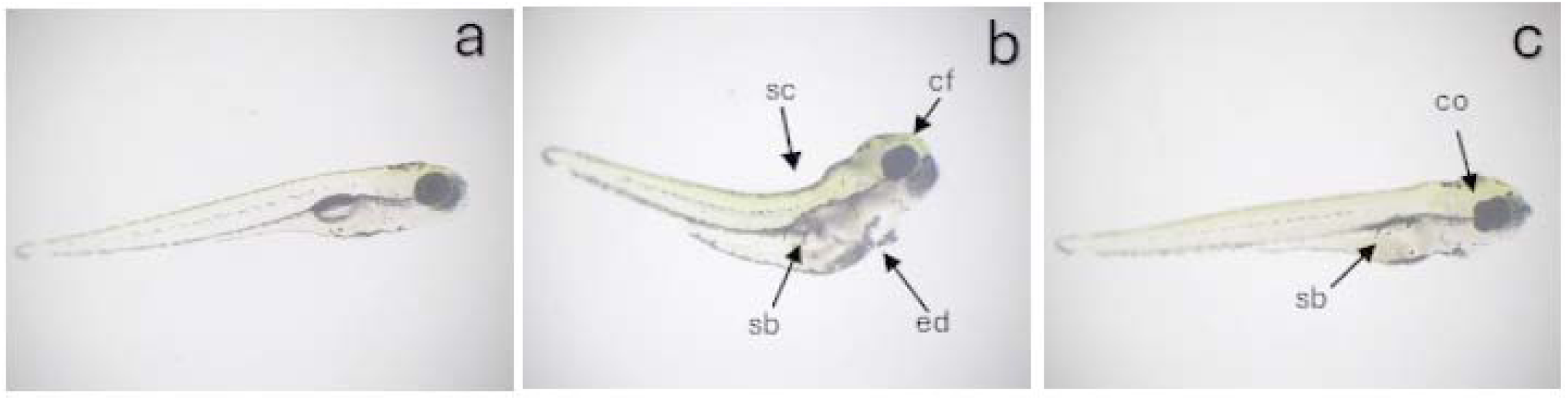
Common phenotypes of developmental toxicity in five-day old zebrafish larvae. Presented above are an unexposed larva showing normal development (a), a larva exposed to 450 µM PFOA (b), and larvae exposed to 80 µM PFHxS (c). Note: cf, craniofacial malformation; sb, swim bladder not inflated; sc, spinal curvature; ed, edema; co, coloboma.

**Figure 3.**
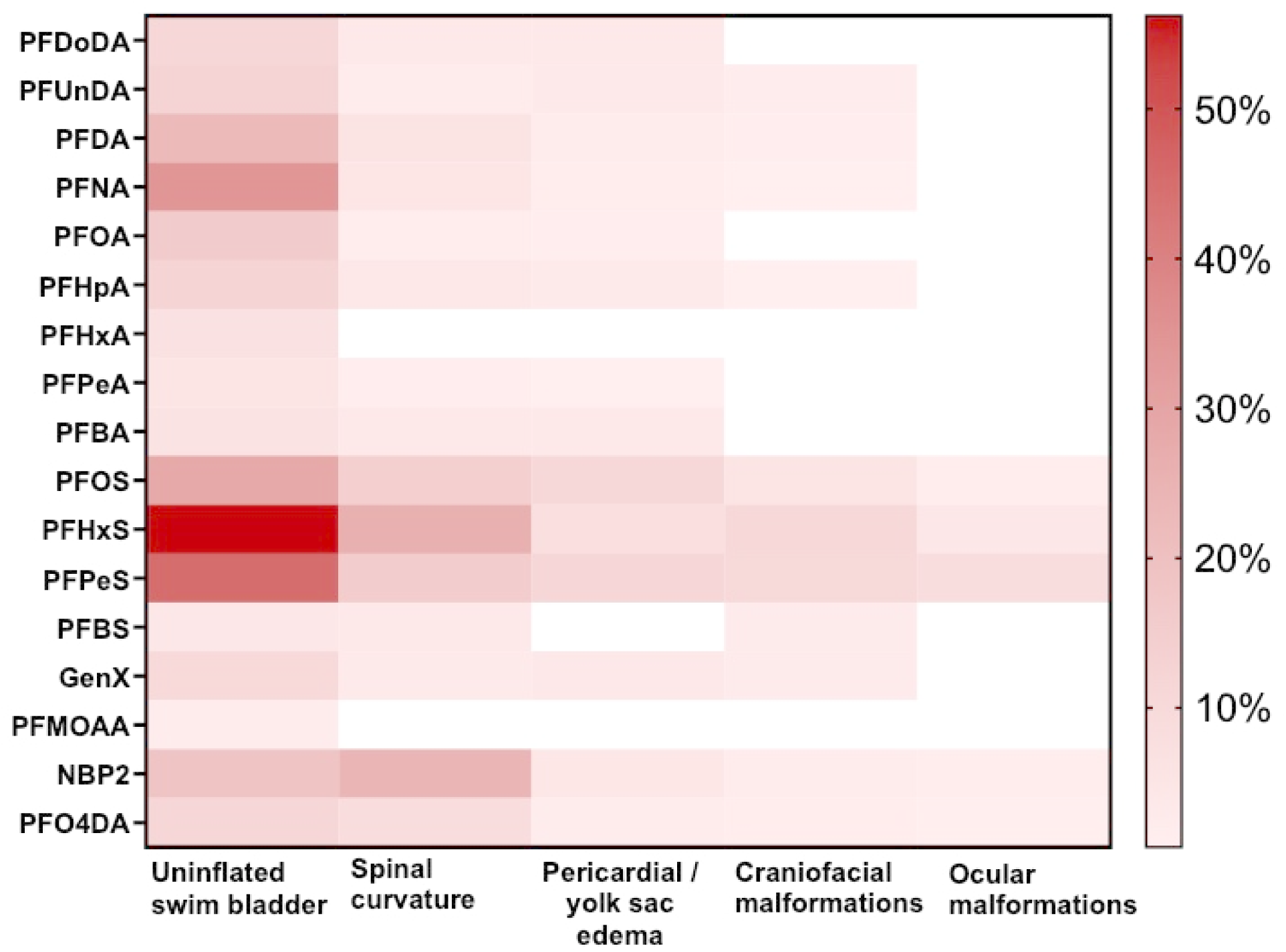
Percent incidence above background for each developmental toxicity endpoint for all exposure wells combined. Any spaces with p > 0.05 were left blank.

**Table 3.** Frequency of failed swim bladder inflation in zebrafish larvae 120 hours post-fertilization following exposure to the specified PFAS. The median effective concentration (EC_50_) is displayed, as well as 20% incidence of failed swim bladder inflation (EC_20_) and 10% incidence (EC_10_). Compounds marked as N/A did not show a dose-dependent response at tested concentrations, and so dose-response curves could not be calculated. PFCA: perfluorocarboxylic acid, PFSA: perfluorosulfonic acid, PFECA: perfluoroethercarboxylic acid, PFESA: perfluoroethersulfonic acid.

|  | Compound | EC<sub>50</sub> [95% CI]<br>(μM) | EC<sub>20</sub> [95% CI]<br>(μM) | EC<sub>10</sub> [95% CI]<br>(μM) |
| --- | --- | --- | --- | --- |
| **PFCAs** | **PFDODA** | 17.52 [14.87-21.08] | 10.34 [7.91-12.89] | 7.59 [5.11-10.24] |
|  | **PFUnDA** | 14.21 [11.75-18.56] | 7.69 [5.43-10.47] | 5.37 [3.10-8.51] |
|  | **PFDA** | 47.69 [37.48-70.4] | 12.38 [8.53-16.62] | 5.63 [2.87-9.24] |
|  | **PFNA** | 61.67 [47.81-82.63] | 20.09 [11.61-29.66] | 10.42 [4.49-18.36] |
|  | **PFOA** | 550 [397.9-816.4] | 181.9 [111.1-276.3] | 95.2 [45.04-170.6] |
|  | **PFHpA** | 3560 [2966-4568] | 2026 [1368-2690] | 1457 [778.2-2180] |
|  | **PFHxA** | 10138 [8945-11233] | 7932 [7011-8926] | 6872 [5755-7936] |
|  | **PFPeA** | N/A | N/A | N/A |
|  | **PFBA** | N/A | N/A | N/A |
| **PFSAs** | **PFOS** | 3.36 [2.56-5.2] | 2.06 [1.06-3.07] | 1.55 [0.52-2.67] |
|  | **PFHxS** | 29.47 [23.01-36.06] | 15.46 [8.78-21.98] | 10.6 [4.78-17.34] |
|  | **PFPeS** | 145.1 [114.7-180.2] | 89.56 [55.99-127.4] | 67.54 [34.88-113.9] |
|  | **PFBS** | 1757 [1218-2740] | 777.2 [469.2-1125] | 482.3 [218.3-?] |
| **PFECA**<br>**s** | **HFPO-DA**<br>**(GenX)** | 4717 [4001-5852] | 2981 [2248-?] | 2279 [1471-?] |
|  | **PFMOAA** | N/A | N/A | N/A |
|  | **PFO4DA** | 1053 [985.5-1124] | 715.5 [639.3-793.9] | 570.9 [487.7-657.6] |
| **PFESA**<br>**s** | **Nafion**<br>**Byproduct 2** | 32.75 [30.6-34.75] | 27.26 [25.77-29.21] | 24.49 [22.81-26.55] |

Spinal malformations were the second most common developmental toxicity phenotype. The majority of spinal malformations were kyphosis (ventral curvature), but lordosis and scoliosis (dorsal and lateral curvature) were also observed. 15 of 17 PFAS caused increased incidence of spinal malformations (Figure 3). Of those, EC_50_ values could be derived for five congeners (PFDoDA, PFOS, PFHxS, PFPeS, NBP2) (Table S1). PFHxA and PFMOAA were the only PFAS that did not cause spinal malformations within the tested range (Figure 3).

The incidence of pericardial and yolk sac edema was significantly elevated across all tested concentrations for 14 PFAS when compared to unexposed controls (Figure 3). Swelling ranged from mild to severe and was highly variable between individuals. PFHxA, PFHxS, and PFMOAA did not cause edema across tested ranges. Only three PFAS (GenX, NBP2, PFO4DA) showed a sufficiently strong dose-response relationship to derive EC_50_s for edema (Table S2).

Craniofacial and ocular malformations were more sporadic. Craniofacial malformations were typically characterized by shortened jaw length and edema, giving the head a smaller, rounded appearance. Fish with this phenotype almost always also had pericardial and yolk sac edema. 11 PFAS exposures resulted in increased craniofacial malformations (Figure 3). Ocular malformations included colobomas and reduced eye size. These malformations were the least common of all assessed phenotypes, elevated in only five exposures (PFOS, PFHxS, PFPeS, NBP2, PFO4DA) (Figure 3).

### Correlations between structure and toxicity

Chain length, head group, and number of ether oxygens were predictive of potency (Figures 4 and 5). Sulfonic acid head groups were consistently more toxic than carboxylic acids. For compounds with equal chain lengths, sulfonic acids without ether oxygens had lower LC_50_ and EC_50_ values in all cases. Four out of five sulfonic acids (PFOS, PFHxS, PFPeS, NBP2) caused ocular malformations, while only one of 12 carboxylic acids did.

**Figure 4.**
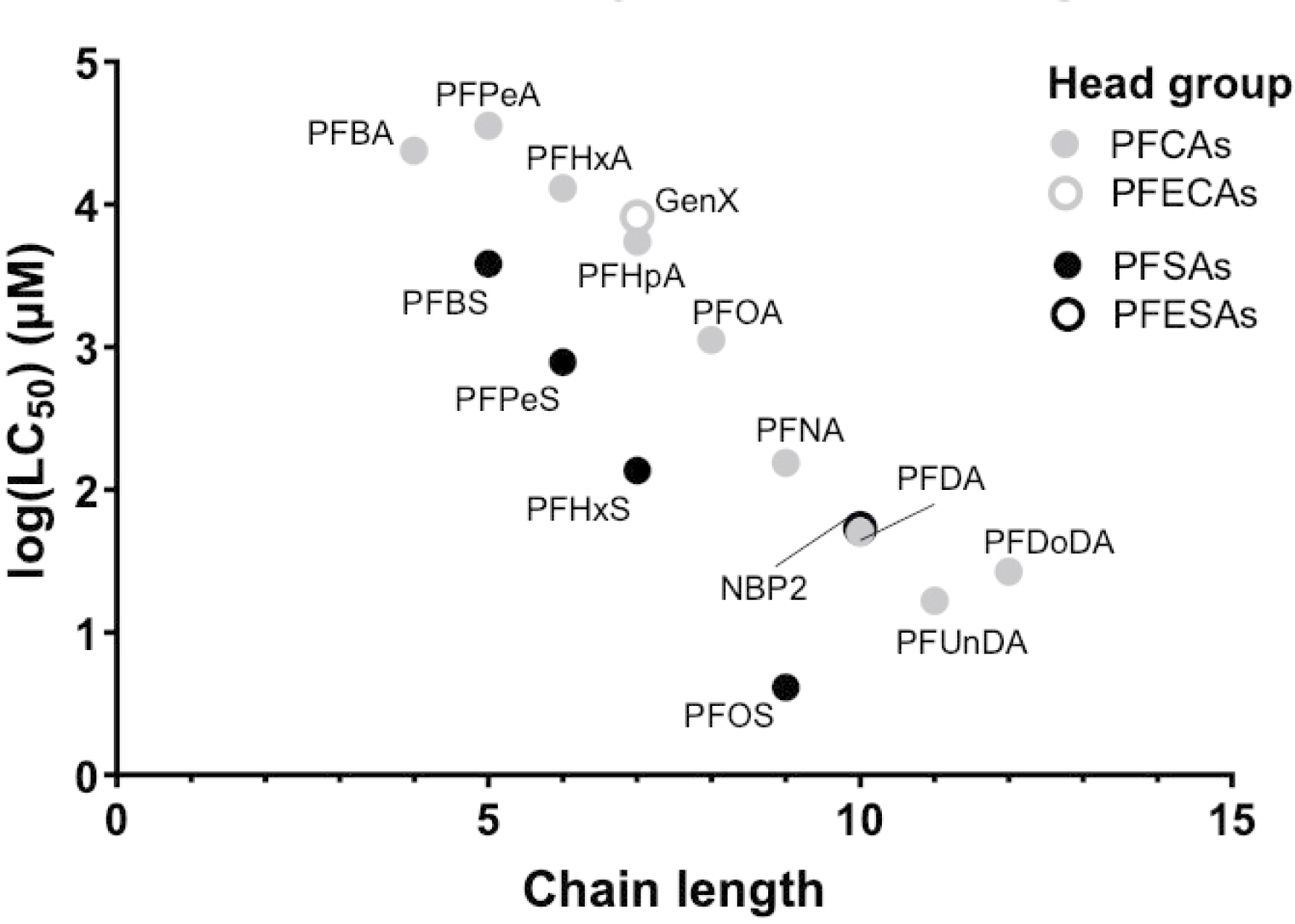
120-hour LC_50_ values compared with PFAS chain lengths. PFO4DA and PFMOAA are excluded because an LC_50_ could not be calculated for either compound.

**Figure 5.**
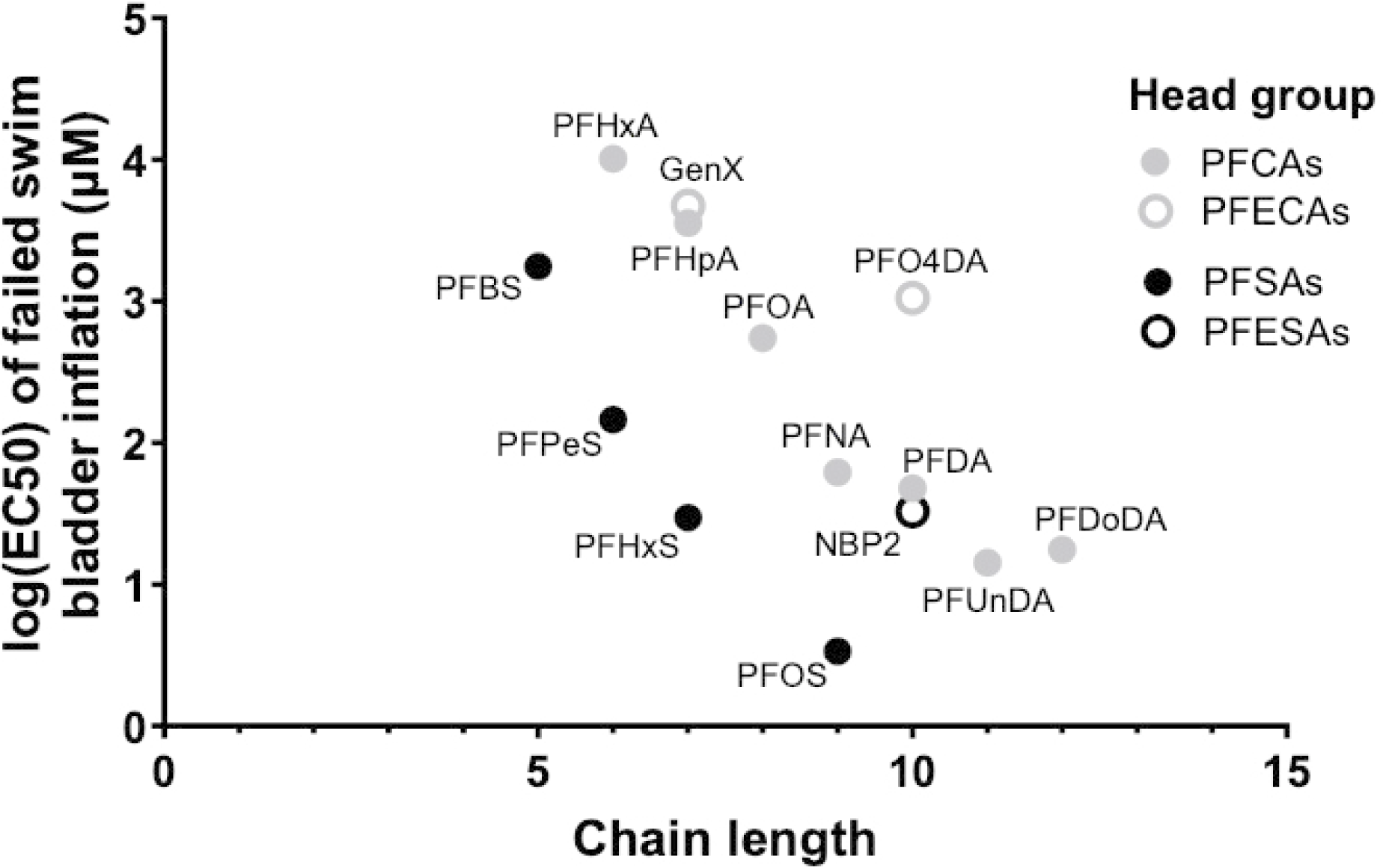
120-hour EC_50_ values for failed swim bladder inflation compared with PFAS chain lengths. PFBA, PFPeA, and PFMOAA are excluded because an EC_50_ could not be calculated for these compounds.

Both lethality and developmental toxicity decreased with the addition of ether oxygens. PFEAs consistently had higher LC_50_s and EC_50_s than PFAAs with the same head group and similar chain length (Figures 4 and 5). LC_50_ values could only be calculated for two PFEAs (GenX and NBP2). However, the other two ether acids in our exposures (PFMOAA and PFO4DA) did not cause elevated mortality at concentrations greater than the LC_50_ of PFCAs with equal chain length, indicating lower potency. We were able to estimate a developmental toxicity EC_50_ for three PFEAs (GenX, NBP2, and PFO4DA), all of which were less potent compared to PFAS without ether linkages.

Other than ocular malformations, none of the endpoints for developmental toxicity showed a significant correlation with any evaluated structural properties. While potency varied greatly between compounds, the developmental impacts of different PFAS were largely similar at toxic concentrations.

Our results could be effectively approximated with MLR models using chain length, head group, and number of oxygens for LC_50_ and EC_50_ of failed swim bladder inflation.

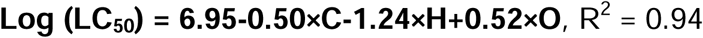

*Equation 1. Multiple linear regression equation for 5-day log (LC_50_) for PFAS in zebrafish. C is chain length, H is the head group (0=carboxylic acid, 1 = sulfonic acid), and O is the number of ether oxygens*.

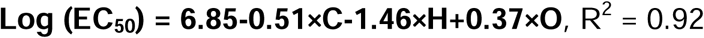

*Equation 2. Multiple linear regression equation for log (EC_50_) of developmental toxicity for PFAS in zebrafish. C is chain length, H is the head group (0=carboxylic acid, 1 = sulfonic acid), and O is the number of ether oxygens*.

Both models aligned closely with our experimental results. Deviation from the LC_50_ model was low for most PFAS but greater in the most (PFOS, PFDoDA) and least (PFBS) potent compounds (Figure S1). Variance in the EC_50_ model was more evenly spread across the tested range (Figure S2). To evaluate the general applicability of these models, we compared predicted LC_50_ and EC_50_ values with experimental values in the literature. Only studies using neutral pH exposure solutions were selected due to the large impact of pH on PFAS toxicity (Wasel et al., 2021). Only a small number of PFAS have published pH-neutral LC_50_ or EC_50_ values in zebrafish, most of which overlap with compounds used in this study. Literature values for four of six PFAS (PFBA, PFOA, PFBS, GenX) were within the 95% CI of our model, although PFBS and GenX were only barely within range (Table 4). Both PFAS outside of the 95% CI had higher published LC_50_ values compared to our prediction. Only PFOA and perfluorotrioxaoctanoic acid (PFO3OA) had published EC_50_ of developmental toxicity values within the 95% CI of our MLR model (Table 5). PFO3OA and perfluoropentaoxadodecanoic acid (PFO5DoDA) were the only compounds not also included in our study.

**Table 4.** A multiple linear regression model was used to predict LC_50_s for select PFAS in zebrafish larvae. These predictions are compared with existing LC_50_ values in the literature. a: Wasel et al., 2022, b: Wasel et al., 2021, c: Wasel et al., 2023, d: Ding et al., 2013, e: Godfrey et al., 2017, f: Hagenaars et al., 2011.

| PFAS | Chain length | Head group | Ether oxygens | Predicted log(LC<sub>50</sub>) [95% CI] (μM) | Literature log(LC<sub>50</sub>) (μM) |
| --- | --- | --- | --- | --- | --- |
| PFBA | 4 | PFCA | 0 | 4.94 [4.51-5.37] | 4.66<sup>b</sup> |
|  |  |  |  |  | 4.81<sup>e</sup> |
| PFHxA | 6 | PFCA | 0 | 3.93 [3.63-4.23] | 4.59<sup>a</sup> |
|  |  |  |  |  | 4.48<sup>b</sup> |
| PFOA | 8 | PFCA | 0 | 2.92 [2.67-3.18] | 3.13<sup>a</sup> |
|  |  |  |  |  | 3.07<sup>b</sup> |
|  |  |  |  |  | 2.95<sup>d</sup> |
|  |  |  |  |  | 3.06<sup>e</sup> |
| PFBS | 5 | PFSA | 0 | 3.19 [2.77-3.62] | 3.62<sup>b</sup> |
| PFOS | 9 | PFSA | 0 | 1.18 [0.75-1.60] | 2.00<sup>d</sup> |
|  |  |  |  |  | 1.75<sup>f</sup> |
| GenX | 7 | PFEC A | 1 | 3.95 [3.49-4.41] | 4.41<sup>c</sup> |

**Table 5.** A multiple linear regression model was used to predict EC_50_ values for select PFAS in zebrafish embryos. These predictions are compared with existing EC_50_ values in the literature. PFO3OA: perfluoro (3,5,7-trioxaoctanoic) acid; CAS 39492-89-2, PFO5DoDA: perfluoro (3,5,7,9,11-pentaoxadodecanoic) acid; CAS 39492-91-6. a: Hagenaars et al., 2011, b: Wang et al., 2020.

| PFAS | Chain length | Head group | Ether oxygens | Predicted log(EC<sub>50</sub>) [95% CI] (μM) | Literature log(EC<sub>50</sub>) (μM) |
| --- | --- | --- | --- | --- | --- |
| PFOA | 8 | PFCA | 0 | 2.78 [2.49-3.08] | 2.39<sup>a</sup> |
|  |  |  |  |  | 2.78<sup>b</sup> |
| PFBS | 5 | PFSA | 0 | 2.85 [2.41-3.30] | 3.7<sup>a</sup> |
| PFOS | 9 | PFSA | 0 | 0.82 [0.40-1.24] | 0.34<sup>a</sup> |
| PFO3OA | 8 | PFEC A | 3 | 3.90 [3.31-4.48] | 3.59<sup>b</sup> |
| PFO4DA | 10 | PFEC A | 4 | 3.25 [2.54-3.97] | 2.49<sup>b</sup> |
| PFO5DoDA | 12 | PFEC A | 5 | 2.61 [1.71-3.51] | 1.72<sup>b</sup> |

## DISCUSSION

Insufficient knowledge of the health risks presented by most PFAS is a serious concern given the ubiquity of these contaminants. Our aim was to provide toxicity data for a range of PFAS, including short-chain compounds and PFEAs. Additionally, we hoped to correlate structural features of the selected PFAS with their toxicity profiles.

Every PFAS that we tested was a developmental toxicant. Similar teratogenic effects were observed across structurally diverse congeners, although potency varied widely. We found that chain length and head group were important structural determinants of potency for PFAAs. Sulfonic acids were more potent than carboxylic acids, and chain length correlated positively with potency. These results are consistent with conventional wisdom on PFAS toxicity, although other studies in zebrafish have not always supported these conclusions. Truong et al. and Britton et al. screened large PFAS libraries (139 and 182 compounds, respectively) for developmental toxicity and did not see significant correlations with chain length or head group. These studies compared a much greater variety of structures, including PFAAs, fluorotelomer alcohols, perfluoroalkyl amides, and silicon PFAS (Truong et al., 2022; Britton et al., 2024). The numerous structural differences between compounds may have obscured the impacts of chain length and head group. In our study, the correlation between chain length and potency was more apparent when viewing carboxylic and sulfonic acids separately (Figures 4 and 5). Chain length and sulfonic acid head groups are also associated with increased bioaccumulation. These PFAS have greater affinity for serum albumin and other proteins, leading to longer half-lives (Rosato et al., 2024). Slower elimination could result in greater internal PFAS concentrations over a five-day exposure period (M. Sun et al., 2022).

The number of ether oxygens was also associated with reduced potency. The inclusion of oxygens in the carbon chain increases solubility, which may reduce bioaccumulative potential by improving renal elimination (Rice et al., 2021) Some support for this can be found in the GenX Exposure Study, which followed a population exposed to PFEAs through drinking water. Short-chain compounds were present at high levels in drinking water but were not detectable in serum samples, while legacy PFAS were widely detected in serum despite lower water concentrations (Kotlarz et al., 2020). Most estimations of PFEA half-lives are shorter than PFAAs of equal chain length, although data are very limited (Wallis et al., 2021). Some PFEAs are unstable in DMSO, and so it was not used in PFEA stock solutions or exposures. DMSO readily permeates biological membranes, increasing absorption of solutes (He et al., 20212). This may have caused increased absorption of PFAAs compared to ether compounds, exacerbating the difference in potency.

Failed swim bladder inflation was the most sensitive developmental toxicity endpoint that we evaluated. Primary inflation of the swim bladder is crucial for swimming performance, and failed inflation is connected to reduced survival (Hagenaars et al., 2014). Thyroid hormones (THs) are crucial for proper inflation of the posterior swim bladder. Disruption of THs results in reduced production of surfactant proteins in zebrafish, leading to swim bladder collapse (Godfrey et al., 2017; Zheng et al., 2011; Horie et al., 2022). Many PFAAs bind effectively to transport proteins in the blood, competitively inhibiting the transport of THs (Ren et al., 2016). Long-chain PFAS and sulfonic acids have a stronger affinity for transthyretin (TTR), potentially explaining the higher potency of those compounds (Dharpure et al., 2022). Failed swim bladder inflation has been observed in exposures to PFAS and other thyroid-disrupting chemicals (Godfrey et al., 2017; Horie et al., 2022). One study showed it was possible to partially rescue PFAS-induced swim bladder defects via supplementation with T3 and T4 (Wang et al., 2020), suggesting supplementation with thyroid hormone may be a strategy to minimize PFAS-mediated thyroid toxicity.

Thyroid disruption may also play a role in other observed developmental toxicity outcomes of PFAS exposure. The role of THs in zebrafish development includes regulation of numerous genes involved in development of the notochord (Shkil et al., 2019). Abnormal curvature of the notochord is commonly reported in PFAS-exposed zebrafish larvae (Albers et al., 2024; Britton et al., 2024). Similar malformations are seen in larval zebrafish exposed to TH inhibitors such as methimazole and phenylthiourea (Chen et al., 2019; Elsalini and Rohr, 2003). Spinal malformations in fish may also occur secondary to failed swim bladder inflation. The impaired swimming that results from an underinflated swim bladder places increased mechanical strain on the spine, leading to malformations (Goolish and Otutake, 1999; Chatain, 1994). An inflated swim bladder exerts some pressure on the developing spine. The absence of this pressure when the swim bladder is not properly inflated can induce abnormal curvature in the spine (Iwasaki et al., 2017).

Ocular malformations were the only endpoint that exhibited a degree of specificity based on PFAS structure. Four of five sulfonic acids (all except PFBS) caused ocular developmental toxicity, while only a single carboxylic acid (PFO4DA) did the same. This is consistent with other studies in the literature, although ocular toxicity is an uncommonly assessed outcome. Several studies have found long-chain sulfonic acid PFAS to induce eye damage and reduced eye size in zebrafish embryos (Lee et al., 2023; Wu et al., 2022). One study comparing structures found that PFOS was far more potent than PFOA at causing reduced eye size, while short-chain compounds did not affect ocular development (Lee et al., 2023). PFAS, particularly long-chain sulfonic acids, are cytotoxic due to their ability to induce oxidative stress by disrupting mitochondrial membrane potential (Tatarczuch et al., 2025; Fang et al., 2010; Hu and Hu, 2009). Apoptosis due to oxidative stress can cause coloboma, reduced eye size, and malformations during ocular development in zebrafish larvae (Kim et al., 2019). Oxidative stress leading to cell death may explain the ability of PFSAs to cause ocular malformations.

Edema is a common response to developmental toxicant exposure in zebrafish, with many possible causes (Wiegand et al., 2023). Damage to the heart and surrounding vasculature due to oxidative stress can cause fluid leakage into the surrounding tissue (Yang et al., 2023). Alterations to cardiac rhythm and output have been observed in conjunction with pericardial and yolk sac edema in PFAS-exposed zebrafish (Yang et al., 2023; Huang et al., 2010). PFAS have also been shown to affect osmoregulation in fish (Lu et al., 2021). Freshwater fish are hyperosmotic relative to their environment, and so reduced osmoregulatory function can result in an excessive influx of water (Hill et al., 2004).

There is considerable variation in the potency of PFAS reported in the literature. Our results are in line with many of these studies (Wasel et al., 2021; Ding et al., 2013; Godfrey et al., 2017; Ulhaq et al., 2013; Hagenaars et al., 2011). Others have found that PFAS are toxic at far lower concentrations than those presented here (Gebreab et al., 2020; Gebreab et al., 2025; Satbhai et al., 2022). Exposure conditions vary widely between different labs, making direct comparisons difficult (Hamm et al., 2019). The pH of exposure solutions is one factor that likely explains some of this variance. Many anionic PFAS are strong acids, but these compounds are often degradation products of larger neutral precursors (Mejia-Avendaño, 2016). We neutralized all exposure solutions to better focus on PFAS-specific toxicity and eliminate pH-dependent outcomes that may not be relevant in real-world exposures. One study comparing the toxicity of buffered and unbuffered PFAS solutions in zebrafish embryos found that the acidic solutions were up to two orders of magnitude more potent than their buffered counterparts (Wasel et al., 2021). In developmental zebrafish, the chorion may also affect PFAS toxicity. Some labs choose to dechorionate embryos prior to exposure, while we left the chorion intact to more closely reflect real-world exposures. The chorion has a limited protective effect against PFAS and other toxicants during the first several days of development (Mylroie et al., 2020). This may affect the ability to compare toxicity data between labs using different protocols.

A model based on chain length, head group, and number of ether oxygens was able to closely approximate our experimental results. This provides support for the feasibility of using structural features to predict PFAS toxicity. The development of structure-based modeling approaches has been a priority to address the large number of distinct PFAS (Sosnowska et al., 2023). Previous studies have used other parameters to create MLR models of PFAS toxicity in zebrafish (Kar et al., 2018; Zhang et al., 2021). Compared with these models, ours had a stronger correlation with testing data, supporting the importance of chain length, head group, and ether oxygens in determining potency.

A significant limitation of our model is the limited number of PFAS in our tests. We examined only PFAAs with sulfonic or carboxylic acid head groups, chain lengths between four and 12, and compounds with 0-4 ether oxygens. Toxicity testing of more diverse structures is needed to provide better data for predictive modeling (Kar et al., 2018). The high variance in toxicity values between studies makes it difficult to create effective models based on multiple datasets (Sosnowska et al., 2023). Even when compared to other zebrafish studies using pH-neutral exposures, there were major differences in experimentally derived LC_50_ and EC_50_ for the same PFAS. This highlights the need for improved standardization of toxicity testing protocols.

## CONCLUSIONS

This study found that structurally diverse PFAS elicit similar phenotypes of developmental toxicity. Failed swim bladder inflation was the most sensitive endpoint for every PFAS tested. The swim bladder is homologous to mammalian lungs, and there is some epidemiological evidence that intrauterine PFAS exposure can impair lung development (Bi et al., 2021; Kung et al., 2021). Potency of PFAS is driven by chain length and head group. The effects of ether linkages on PFAS toxicity were not conclusive. A unique strength of this study is the comparison of better understood legacy PFAS with short-chain alternatives and PFEAs in a neutral pH environment. The anionic PFAS used in many studies are strong acids, and toxicity due to low pH can overshadow chemical-specific toxicity. We believe that the results of this study more accurately reflect the impacts of key structural properties on PFAS toxicity.

There is still a need for further toxicity testing on most PFAS. The inclusion of more PFEAs in future studies is necessary to understand the impact of ether substitutions. A better understanding of the toxicokinetics of PFAS is another crucial goal to elucidate how aqueous concentrations connect to internal dosages. Improved knowledge of structural properties on various aspects of toxicity should be incorporated into predictive models, allowing for identification of less toxic alternatives to the PFAS currently in use.

## AUTHOR CONTRIBUTIONS

Conceptualization: MF and AP; sample collection: RB and MF; analysis: RB, MF, EG, AP; original draft write-up: MF; original draft edit and review: AP; project management: AP; funding acquisition: AP. All authors have read and agreed to the submitted version of the manuscript.

## ACKNOWLEDGEMENTS

We thank members of the Planchart lab for valuable discussions during the execution of this work.

